# Cellular Pharmacology of Curcumin With and Without Piperine

**DOI:** 10.1101/2021.07.14.452424

**Authors:** Rama I Mahran, Pan Shu, Justin Colacino, Magda M Hagras, Duxin Sun, Dean E Brenner

**Author notes:** ***Correspondance:*** Dean E Brenner, 2150 Cancer Center, University of Michigan Medical Center, Ann Arbor, MI 48109-5290.

## Abstract

Prior reports have suggested that piperine enhances curcumin’s anti-carcinogenesis. We tested the hypothesis that piperine increases the intracellular concentrations of curcumin by improving intracellular uptake or reducing curcumin efflux or metabolism in breast cells. We incubated SUM149, MCF10A, primary normal human breast cells, ALDH^+^, and ALDH^-^CD44^+^24^-^ SUM149 cells with curcumin ± piperine at concentrations 1 μM to 15 μM for time periods of 15 minutes to 24 hours. We assayed cell viability by MTT assay and proliferation by primary mammosphere assay. Curcumin and its metabolites were assayed using liquid chromatography mass spectroscopy. Curcumin, but not piperine, showed significantly higher effects on the viability of breast cancer SUM149 cells than in non-tumorigenic MCF10A cells. Curcumin + piperine synergistically reduced viability of SUM149 cells but had a concentration dependent effect upon MCF10A cell viability. Cellular uptake of curcumin in SUM149 is significantly higher, while the efflux in SUM149 is significantly lower than in MCF10A, which correlated with cell viability. Piperine did not alter curcumin cellular uptake, efflux, or metabolism in any of the cell models. The observed synergism of piperine+curcumin in reducing breast stem cell self renewal is likely due to independent anti-carcinogenesis effects rather than any effects upon intracellular curcumin concentrations.

## Introduction

Curcumin—the major bioactive product extract from the rhizome of the *Curcuma longa* plant (turmeric)— has a wide spectrum of anti-carcinogenic, antioxidant (1), anti-inflammatory (1), and cancer chemopreventive effects in colon cancer (2), breast cancer (3), and other types of cancers (4). The molecular mechanisms by which curcumin exerts these broad biological effects are diverse and include inhibition of eicosanoid synthesis (5, 6), inhibition of NFκB release (5, 6), inhibition of stemness through interruption of Wnt signaling (5, 7, 8), modification of cellular lipid profiles (9), modification of P-glycoprotein expression (10), and inhibition of vascular endothelial growth factor (VEGF)-mediated angiogenesis (5, 6). This broad group of anti-carcinogenesis mechanisms has translated into potent chemopreventive effects in classical chemical *in vivo* rodent carcinogenesis models of diverse organ systems (3, 11-14) as well as in transgenic rodent carcinogenesis systems (3, 14-16).

Despite a broad anti-carcinogenesis effect in preclinical modes, curcumin has not found a role in clinical cancer prevention, primarily due to poor bioavailability in humans. To address this barrier, investigators have sought to enhance absorption and systemic bioavailability by inhibiting cell membrane efflux transporters, reducing metabolism and conjugation by inhibiting Phase I and II metabolizing enzymes by using bioenhancers (4, 17), creating novel delivery systems to overcome first pass extraction and rapid systemic metabolism (4, 18), and by synthesizing analogs or conjugates to enhance curcumin’s solubility and stability while retaining its pharmacological properties.

Piperine, an alkaloid extract from pepper, has been studied *in vivo* as a bioenhancer of curcumin because of its inhibitory properties on cellular membrane efflux transporters (19-21) and phase I and phase II metabolizing systems (20, 22-26). An early clinical trial of piperine combined with curcumin (17) suggested that piperine enhanced curcumin’s systemic bioavailability. More recent data using contemporary analytical technologies and Good Manufacturing Practices compliant formulations have failed to reproduce these initial findings (27-29).

We found that piperine alone inhibits normal human breast mammosphere formation (7, 30). In combination with curcumin, piperine enhanced curcumin inhibition of primary normal human breast epithelial ALDH^+^ cells via inhibition of the Wnt signaling pathway (7). These data suggested that piperine’s interaction with curcumin may be independent of its bioenhancer activity. Data in cell culture systems support piperine’s anti-carcinogenic activity. For example, piperine inhibits activator protein 1 (AP-1) and nuclear factor-κB (NF-κB) through the inhibition of ERK1/2, p38 MAPK, and Akt signaling pathways in SKBR breast cancer cells (31). Piperine inhibits Wnt **β**-catenin signaling pathway in colorectal cancer cells (32).

The issue of whether piperine functions as a bioenhancer of curcumin’s anti-carcinogenic activity or as an independent anti-carcinogen with additive or synergistic activity with curcumin remains unresolved. Here we addressed the question of whether piperine’s enhancement of curcumin’s anti-carcinogenic activity is due to enhancement of piperine’s cellular uptake of curcumin or is due to independent, additive or synergistic anti-carcinogenesis effect.

## Materials and Methods

### Chemicals

Curcumin (>90% pure) was purchased from Cayman (Ann Arbor, MI) and piperine (>99% pure) was purchased from Sigma Aldrich (St. Louis, MO). Stock solutions of curcumin and piperine were prepared in dimethyl sulfoxide (DMSO), prepared to a concentration of 50 mM and stored in −80 °C. Curcumin O-sulfate (COS >95%, w/w) was purchased from Toronto Research Chemicals (Toronto, Ontario) and tetrahydrocurcumin was purchased from Santa Cruz Biotechnology (Dallas, TX). The internal standard CE302 (>98%, w/w) was synthesized in the University of Michigan Pharmacokinetic Core lab and was used without further purification. HPLC grade acetonitrile and formic acid were obtained from Sigma Aldrich; Methanol was purchased from Thermo Fisher Scientific (Waltham, MA). HPLC grade water was obtained via a Milli-Q Integral Water Purification System (Darmstadt, Germany).

### Cell Lines

The MCF10A human breast cell line was purchased from ATCC (Manassas, VA). SUM149 was purchased from Asterand (Detroit, MI). MCF10A was cultured in DMEM/F12 lonza (Walkersville, MD) supplemented with 5% horse serum, insulin (10 μg/mL), epidermal growth factor (20 ng/mL), cholera toxin (100 ng/ml), and gentamycin (50 ng/mL). SUM149 cells were cultured in DMEM/F12 Lonza supplemented with 5% fetal bovine serum, insulin (5 μg/mL), and hydrocortisone (1mg/mL).

### Normal Human Breast Mammosphere Assay

#### Normal Human Breast Tissue Dissociation

Normal human breast tissue was obtained from 15 women who were undergoing elective reduction mammoplasty. After providing informed consent (following the University of Michigan IRBMED approved tissue collection protocol), the tissue was obtained through the Tissue Procurement Service with the Department of Pathology at the University of Michigan Medical School. The breast tissue of 15 patients was mechanically and enzymatically dissociated into single cells as previously described (33).

#### Mammosphere Formation

Pooled breast epithelial single cells from 15 patients were cultured in MammoCult media (Stem Cell Technologies, Vancouver, Canada) and treated with curcumin, piperine, or both or the DMSO control (0.1%) in triplicates/dose/combination. Cells were plated in 6-well ultralow attachment plates (Corning Inc, Corning NY) at a density of 100,000 cells/mL. Primary mammospheres were allowed to form for 10 days. The mammospheres’ size was measured using Spot Imaging Software. The mammospheres that were 40 μm or larger were counted manually by two independent observers. The average number and size of mammospheres for each treatment were calculated. The experiment was repeated three times.

### Cell Viability Assay

MCF10A and SUM149 cells were plated at a density of 3×10^3^ cells/well in 96 well plates. The next day, the cells were treated with increasing concentrations of curcumin (0-20 μM) or piperine (0-200 μM) or both or the DMSO control (0.1%) in 6 replicates for each dose or combination, and the cells were incubated for 72 hours. A MTT assay (American Type Culture Collection (ATCC)) was performed per the manufacturer’s instructions. The absorbance of the colored solution was measured using the plate reader (Synergy multimodal reader) at 570 nm wavelength.

### Stability of Curcumin ± Piperine in Complete media and PBS

To determine the type of solvent that would be used in the uptake experiments, we pipetted curcumin into complete media (containing 5% FBS), MammoCult media, or PBS to reach a concentration of 15 μM (the highest curcumin concentration used in the uptake experiments), and the solutions were vortexed well. The solutions were incubated and protected from light at 37 °C (to mimic the cellular incubation conditions) for 0, 0.08, 0.16, 1, 4, and 8 hours. At each time point, the solutions were vortexed; curcumin and its degradation products were extracted and analyzed by the LC-MS system.

To determine the effect of piperine on curcumin stability, solutions of 15 μM curcumin ± 10 μM piperine were prepared in PBS and incubated at 37°C protected from light for 0, 5, 10, and 60 minutes. At each time point, the solutions were vortexed; curcumin was extracted from each solution and quantified by LC-MS. The experiment was repeated three times.

### Cellular Uptake, Efflux and Metabolism of Curcumin ± Piperine

Our data (Figure 1) and those of others (34, 35) show that curcumin is unstable in the physiological buffer. Although MammoCult media is considered serum free media, the stability of curcumin in this type of media was comparable to complete media. *In vitro* experiments were performed in complete media (containing 5% serum).

**Figure 1.**
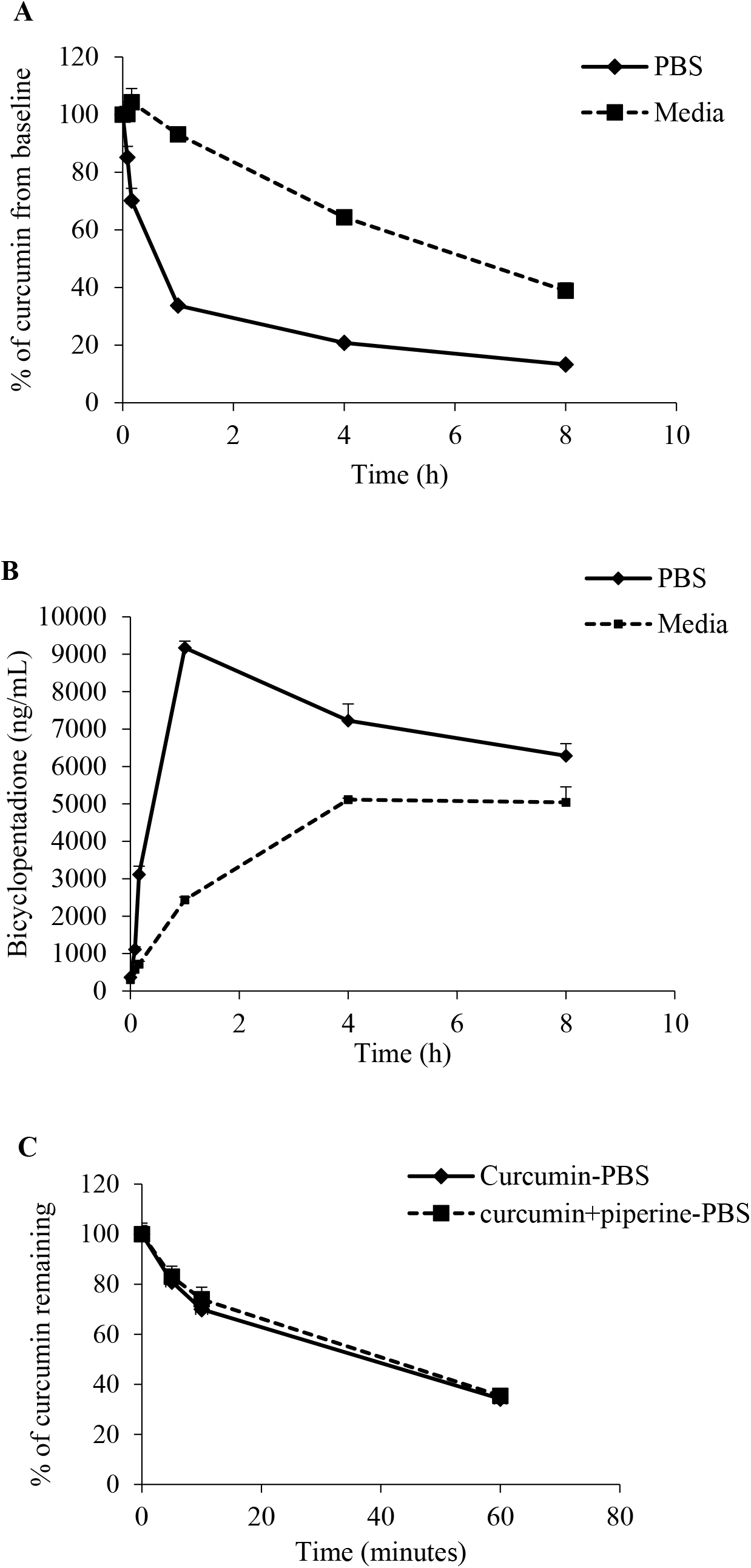
Curcumin is unstable in physiologic solution. A. Curcumin was incubated in either phosphate buffered saline (PBS) or cell media (DMEM/F12 Lonza supplemented with 5% fetal bovine serum, insulin, hydrocortisone) protected from light at 37° C. Samples were then extracted and assayed by LC-MS. Data presented as the percent of curcumin concentration detected to the curcumin concentration at time zero. Within 8 hours, 50% of curcumin is degraded to degradation products. B. Curcumin’s primary degradation product, bicyclopentadione increases in concentration over time in both PBS and media commiserate with the decline in curcumin. C. Piperine does not enhance curcumin stability in PBS. Piperine 10 μM was added to curcumin in the same conditions as shown in panel A. No difference in degradation rate of curcumin was found.

#### Cellular Uptake

SUM149 and MCF10A cells were seeded at a density of 3×10^4^ cells/well in 96 well plates and left to attach overnight. The following day, the cells were washed and incubated with warm HBSS for 2 minutes. The HBSS was then replaced by 15 μM curcumin ± 10 μM piperine for 30, 60, 120, 240, and 360 min, 1 μM curcumin ± 1 μM piperine, or 5 μM curcumin ± 5 μM piperine for 5, 10 and 60 minutes or the DMSO control in complete media alone for 24 hours. Each dose or combination was performed in triplicates (triplicates/dose/combination). At each time point, the media was removed, extracted by methanol and kept at −80 °C for assay of curcumin, metabolites, and degradation products using liquid chromatography-mass spectroscopy (LC-MS). The cell monolayer was washed with cold HBSS three times and lysed by adding 50 μL/well of 1X radioimmuno-precipitation assay (RIPA) cell lysis buffer (Santa Cruz Biotechnology). The plates were maintained at constant agitation at 4°C for 20 minutes, and the cell lysates were collected and sonicated while on ice for 15 seconds using medium power. The extract was assayed by the LC-MS. Protein concentrations in the cell lysates (mg/mL) were assayed using the Bradford protein assay (Bio-Rad, Hercules, CA) per the manufacturer’s instructions. Curcumin concentrations were normalized to protein concentration. To assess whether pre-treatment with piperine might alter the cellular uptake of curcumin, MCF10A cells were plated at a density of 3×10^4^ cells/well in 96 well plates and incubated overnight to attach. The following day, the cells were washed with HBSS, incubated with increasing concentrations of piperine (0-50 μM) in warm HBSS, and incubated for 30 minutes. After 30 minutes, HBSS was replaced by 15 μM curcumin dissolved in complete media, and the cells were incubated for 2 hours. After 2 hours, the media was removed, the cell monolayer washed, lysed, and extracted to be analyzed by LC-MS.

To understand cellular uptake of curcumin in the presence and absence of piperine in a model system where we observed significant effects in primary breast cells, we used pooled primary normal human breast cells obtained from 15 patients undergoing mammoplasty procedures. Pooled cells were seeded at the density of 2×10^5^ cells/well in 24-well ultralow attachment plates in MammoCult media, treated with 15 μM curcumin ± 10 μM piperine or DMSO control in triplicate/dose/combination and incubated for 0.5, 1, 2,4, and 6 hours. At each time point, the cells were collected and washed twice with cold HBSS. The cell pellet was lysed by 100 μL RIPA lysis buffer, and then proceeded with the LC-MS analysis.

#### Cellular Efflux

We incubated SUM149 cells and MCF10A cells (in 2D culture) with 15 μM curcumin ± 10 μM piperine for 1 hour, then the media containing the drugs was removed, the cell monolayer was washed twice, then incubated with fresh media with or without 10 μM piperine for 0.08, 0.25, 0.5, 1, 2, and 4 hours. At each time point, the media was removed and extracted, the cells were lysed and extracted and proceeded with the LC-MS analysis as described above.

#### Assessing Curcumin Uptake in Breast Stem Cells

There are two major populations of breast stem cells: Cells that carry the surface marker of CD44^+^CD24^-^and others which express aldehyde dehydrogenase (ALDH^+^) (36). We purified ALDH^+^ and CD44^+^CD24^-^ cells as well as the non-stem cells from SUM149 cells using MoFlo Astrios flow cytometer (FACS) as described previously (30). Single SUM149 cells were stained by Alexafluor750 LIVE/DEAD Fixable Dead Cell Stain (Life Technologies, Carlsbad, CA), CD24-Brilliant Violet 421 (Biolegend, San Diego, CA), CD44-APC (Biolegend), and Aldefluor (Stem cell Technologies). For compensation and gating purposes, single stain and isotype controls were included. ALDH^+^, CD44^+^CD24^-^, ALDH^-^, CD44^+^/^-^CD24^+^ were sorted by FACS. ALDH^+^ and ALDH^-^CD44^+^CD24^-^ cell populations, and the remaining SUM149 cells that didn’t express any of these markers (non-stem cells) were seeded at a density of 3×10^4^ cells/well in 96 well plates and incubated overnight. The next day, the cells were treated with 15 μM curcumin ± 10 μM piperine for one hour, then washed, lysed, and extracted for LC-MS analysis.

### Liquid Chromatography-Mass Spectroscopy (LC-MS) Assay for Curcuminoids, Metabolites and Degradation Products

We used a LC-MS method to separate and directly quantify curcuminoids (curcumin, demethoxycurcumin, and bisdemethoxycurcumin), curcumin metabolites (curcumin sulfate conjugates, and tetrahydrocurcumin), curcumin degradation products (bicyclopentadione and ferulic acid), and internal standard 7-(3,5-dimethylisoxazol-4-yl)-6-methoxy-2-methyl-N-(1-methyl-1H-pyrazolo[3,4-b]pyridin-3-yl)-9H-pyrimido[4,5-b]indol-4-amine (CE302).

#### Sample Preparation, Extraction and Analyte Stability

Stock solutions of curcumin was prepared in DMSO at a concentration of 50 mM and stored at −80 °C. To prepare a calibration curve, an aliquot of 15 μL of diluted standard solution in methanol (a mixture of curcumin, tetrahydrocurcumin, and curcumin sulfate each 1-5000 ng/mL) was spiked into an aliquot of 15 μL blank cell lysate and media to yield cell lysate and media samples containing 1-5000 ng/mL standards, respectively. The lysates or media (15 μL) was mixed with 15 μL of methanol. These samples were extracted with 100 μL of internal standard solution dissolved in methanol, by vortexing and centrifuging at 2000 x g for 10 min. at 4 °C. An aliquot of 5 μL supernatant was used for the LC-MS/MS analysis.

There is no available bicyclopentadione standard, therefore, we quantified the bicyclopentadione against the standard curve of curcumin sulfate, as described (37). We performed curcumin stability experiments by comparing peak size, area, and fragmentation of curcumin assayed from 15 μM stock solution in either complete media containing 5% fetal bovine serum or standard phosphate buffered saline kept at 37°C for 0, 5, 10, 60, 240, and 480 minutes protected from light. Solutions of 15 μM curcumin ± 10 μM piperine were prepared in PBS and incubated at 37 °C protected from light for 0, 5, 10, and 60 minutes.

#### Equipment

LC-MS was performed on a Shimadzu HPLC system (Shimadzu) consisting of a SCL-20A system controller, two LC-20AD pumps, and an SIL-20AD auto-sampler. Chromatographic peaks were monitored through an API 5500 Qtrap mass spectrometer equipped with an electrospray ionization quadrupole mass analyzer (AB Sciex LLC, Framingham, MA). Analyst software (Version 1.6.2) was used for system control and data processing.

#### Analytical Methodology

The analytes were separated in a mobile phase consisting of two components: mobile phase A, 0.1% formic acid in water and mobile phase B, 0.1% formic acid in acetonitrile. The gradient profile was 0–30 min, 85–0% mobile phase A linear; the flow rate was 0.7 mL/min. The optimized system parameters were obtained via direct infusion of standard solutions, respectively. The ion transition pairs for the quantification of curcumin were selected based on the highest multiple reaction monitoring (MRM) signal under positive (+) mode. The screening and identification of curcumin metabolites were carried out under negative (-) mode for better ionization.

### Data Analysis

The percent of curcumin concentration in complete media and PBS at each time point relative to its concentration at time zero was calculated as follows: (The peak area ratio of curcumin to the IS peak area in the solution at specific time point/ the peak area ratio of curcumin to the IS peak area at time zero) x 100%. To calculate the half-life of curcumin in complete media and in PBS, we added exponential trendlines to the generated graphs in Microsoft Excel and the equations generated from these trendlines were used to estimate that half-lives.

To calculate the percent of cell viability in the MTT assays, the average absorbance of each treatment was divided by the average absorbance of the DMSO control and was then multiplied by 100%.

To determine the nature of the interaction between curcumin and piperine, the combination index (CI) theorem based on the Median-Effect Equation (38) was used. Combination indices were calculated using the Compusyn software (ComboSyn Inc.), and r ≥ 0.95 was considered a simulation for the dose-effect relationship. A CI between 0 and 0.9 indicates synergy; a CI between 0.9 and 1.1 indicates an additive effect, and a CI with a value from 1.1 to infinity indicates an antagonistic interaction (39-41). The IC_50_ of curcumin and piperine were also calculated by Compusyn software. To calculate the ratio of curcumin mass detected in the media to its mass intracellular, we calculated the mass of curcumin in the cell lysates and in the media by multiplying the concentration of curcumin (ng/mL) by the volume of the cell lysates (0.05 mL), or the volume of media (0.1 mL). Then, we divided the mass of curcumin in the media by mass of curcumin in the cells.

Data are expressed as mean ± standard deviation. The following data were compared between groups using Student’s T test: IC_50_ of curcumin and piperine in MCF10A versus SUM149 and mammosphere formation, and the intracellular curcumin concentrations at each time point. The intracellular curcumin concentration, curcumin and its metabolites in the media of the cell models studied were compared between groups using a one-way analysis of variance (ANOVA) using SPSS 17.0 with adjustment to multiple comparison with the Bonferroni method. A p value < 0.05 was considered significant.

## Results

### Piperine Does Not Enhance Curcumin Stability

One potential explanation for piperine enhancement of curcumin anti-carcinogenic effect might be that piperine stabilizes the curcumin molecule in physiologic conditions. Curcumin degrades rapidly in PBS reaching 50% of its original concentration after 0.62 ± 0.002 hours (Figure 1A). In complete cell culture media, curcumin is stable for 1 hour with a slower degradation rate over time than in PBS. Curcumin’s half-life in complete media is 6.53 ± 0.46 hours. The decrease of curcumin concentration over time in the PBS and in complete cell culture media was accompanied by an increase of curcumin’s major degradation products, especially bicyclopentadione (Figure 1B) and ferulic acid (data not shown). We found bicyclopentadione and ferulic acid intracellularly and in the incubation cell culture media (Data not shown). Addition of piperine to curcumin in PBS did not protect curcumin from degradation (Figure 1C).

### Effects Upon Cell Viability of Invasive and Non-Invasive Cell Lines

We measured the effects of a 72 hour treatment of curcumin (0-20 μM) and piperine (0-200 μM) on cell viability in SUM149 and MCF10A cells, comparing the effects to the DMSO controls. The IC_50_ was 8.75 ± 0.15 μM (mean ±standard deviation) (Figure 2A) and 94.8 ± 5.3 μM (Supplemental Figure 1) for curcumin and for piperine (p value of IC_50_ of curcumin compared to piperine<0.001) respectively in SUM149 cells. In SUM149 cells, the combination index (CI) analysis provides evidence of synergism at most curcumin concentrations (Figure 2B). Higher curcumin concentrations (≥20 μM) alone are cytotoxic, thus no synergism can be assessed at those concentrations. The IC_50_ was 15 ± 0.4 μM (Figure 2C) and 114 ±14.3 μM (Supplemental Figure 1) for curcumin and piperine, respectively in the MCF10A cells (p<0.001). While MCF10A cells are more resistant to the cytotoxic effects of curcumin compared to SUM149 (p value of the IC_50_ of curcumin’s cytotoxicity in MCF10A compared to SUM149= 0.004). Combination index analysis suggests additive effects of piperine to curcumin’s effects in MCF10A cells (Figure 2D). Curcumin—piperine cytotoxic interaction is less pronounced in the non-invasive MCF10A cells than in SUM149 cells (Figure 2B and D) with synergy demonstrated only between curcumin and piperine at the lower concentrations of both drugs.

**Figure 2.**
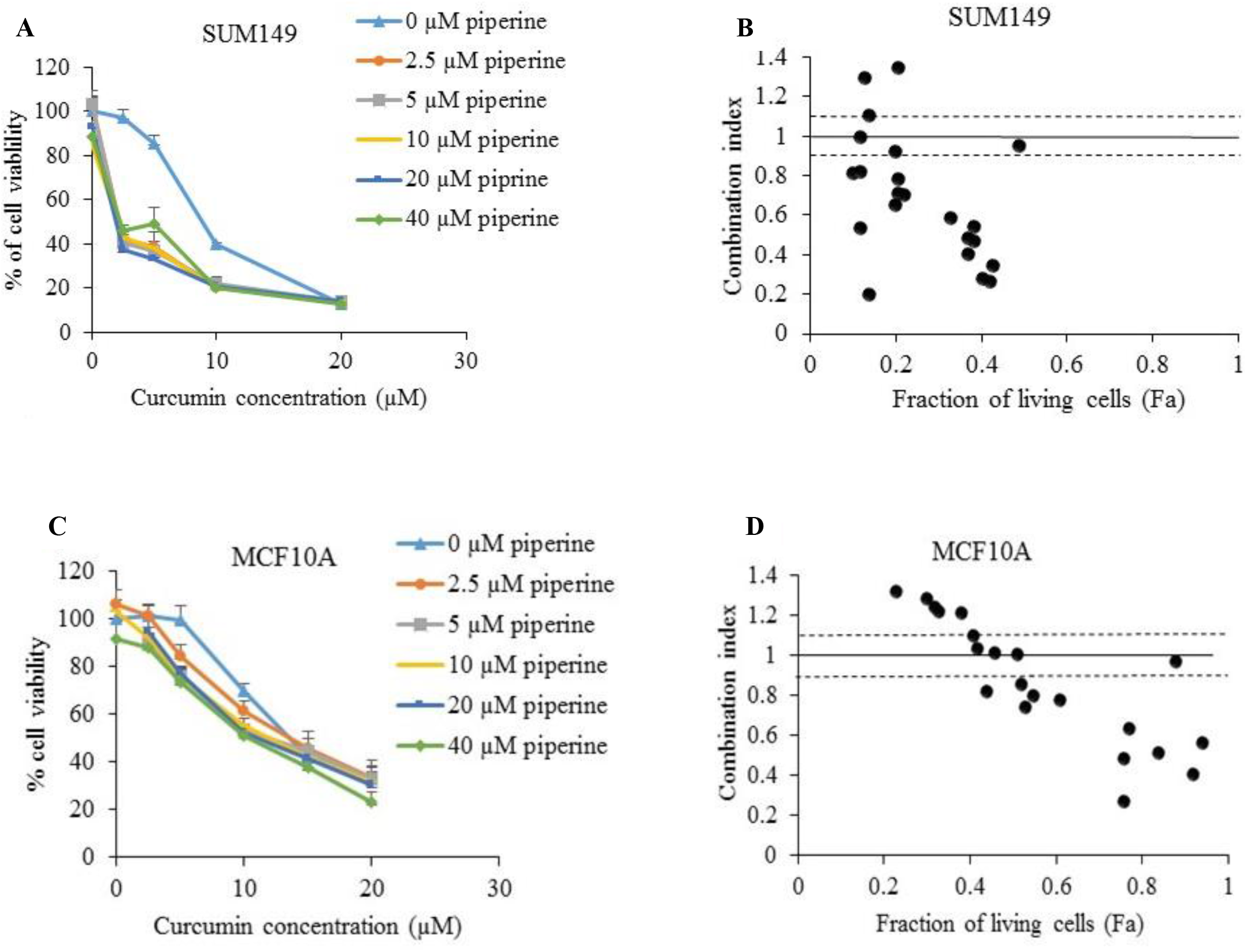
Curcumin ± piperine effects on viability of invasive and non-invasive neoplastic breast cells. Percent of viable MCF10A (A) and SUM149 cells (B) after treatment with curcumin and piperine compared to controls treated with the DMSO vehicle (0.1%) for 72 hours. Results expressed as the mean ± SD of two independent experiments 6 replicates each. Combination index (CI) plots for the effect of combining curcumin and piperine on the fraction of living SUM149 (B) and MCF10A (C) cells. Two points in Panel B showing antagonism were curcumin concentrations of 20 μM, a cytotoxic concentration. CI value < 0.9 indicates synergy, between 0.9-1.1 indicates additive effects and >1.1 indicates antagonistic effects.

### Effects Upon Primary Human Mammosphere Formation

We repeated and verified our previously published data (7) that curcumin and piperine additively inhibits normal human breast stem cell self-renewal using a mammosphere assay as a surrogate for breast stem cell self-renewal (Supplemental Figure 2).

### Cellular Uptake

Previous report suggested that piperine acts as a bioenhancer of curcumin by increasing its intracellular concentration (17). We found no significant increase in curcumin uptake with piperine incubation in MCF10A cells, SUM149 cells, or in the primary normal human breast cells after six hours of incubation with curcumin 15 μM ± piperine 10 μM (Figure 3A) or curcumin 5 μM ± piperine 5 μM (SUM149 and MCF10A only) (Figure 3B). Preincubating MCF10A breast cells with increasing concentrations of piperine (from 1 μM-50 μM) for 30 minutes prior to the addition of 15 μM curcumin for 2 hours did not increase curcumin’s cellular uptake (Data not shown).

**Figure 3.**
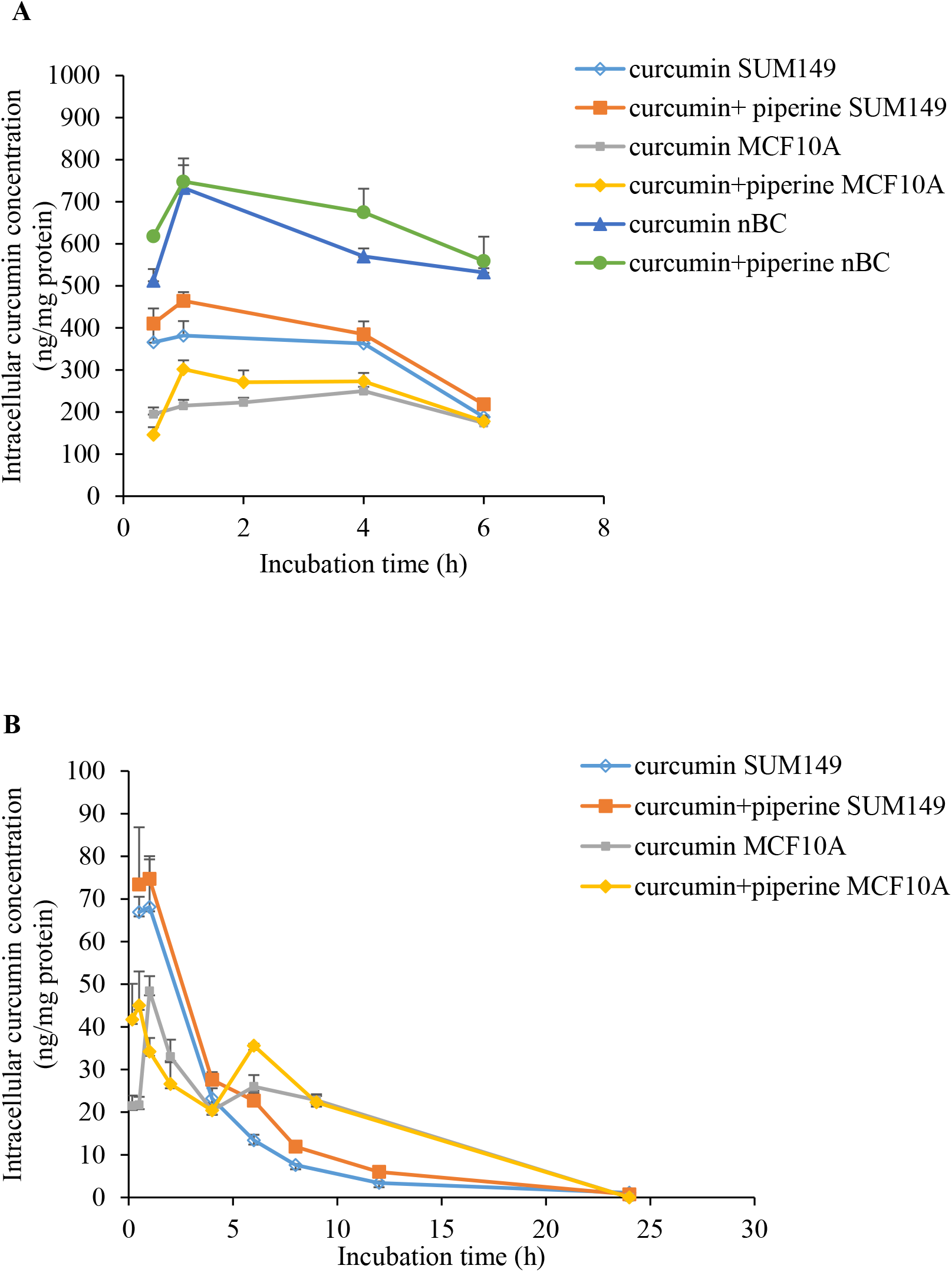
Cellular uptake of curcumin is not altered by piperine. Panel A: Intracellular curcumin concentration after incubating MCF10A, SUM149, and normal human breast cells (nBC) with 15 μM curcumin ± 10 μM piperine for 6 hrs. Panel B: Intracellular curcumin concentration after incubating MCF10A, SUM149 with 5 μM curcumin ± 5 μM piperine for 24 hrs. Results are expressed as the mean ± SD from triplicates. nBC= normal human breast cells.

Intracellular concentrations of curcumin in SUM149 cells were significantly higher than MCF10A cells (Figure 3). For example, after 1 hour of incubation with 5 μM curcumin, the intracellular concentration of curcumin was 68.1 ± 22.3 ng/mg protein and 21.7 ± 3.4 ng/mg protein for SUM149 and MCF10A respectively (p= 0.03). The SUM149 curcumin intracellular concentration after 1 hour is three-fold higher than the MCF10A concentration, while the curcumin IC_50_ in SUM149 cells is two-fold lower than the IC_50_ in MCF10A cells. Differential cellular uptake of curcumin in the invasive breast cell line (SUM149) compared to the non-invasive cell line (MCF 10A) partially explains the differences in IC_50_ observed. We could not compare the intracellular curcumin concentration in primary normal human breast cells to the cell lines since they were cultured in different culture conditions (3D cultures).

Using flow cytometry, we isolated stem cell pools (ALDH^+^, CD44^+^CD24^-^) from SUM149 cells and incubated each of these cell fractions with 15 μM curcumin ± 10 μM piperine for 2 hours. We also incubated the remaining SUM149 that did not express stemness surface markers with the same curcumin ± piperine conditions. We found no difference in intracellular curcumin concentrations in any of the stem cell fractions or in the remaining non stem SUM149 cells with piperine incubation (Figure 4).

**Figure 4.**
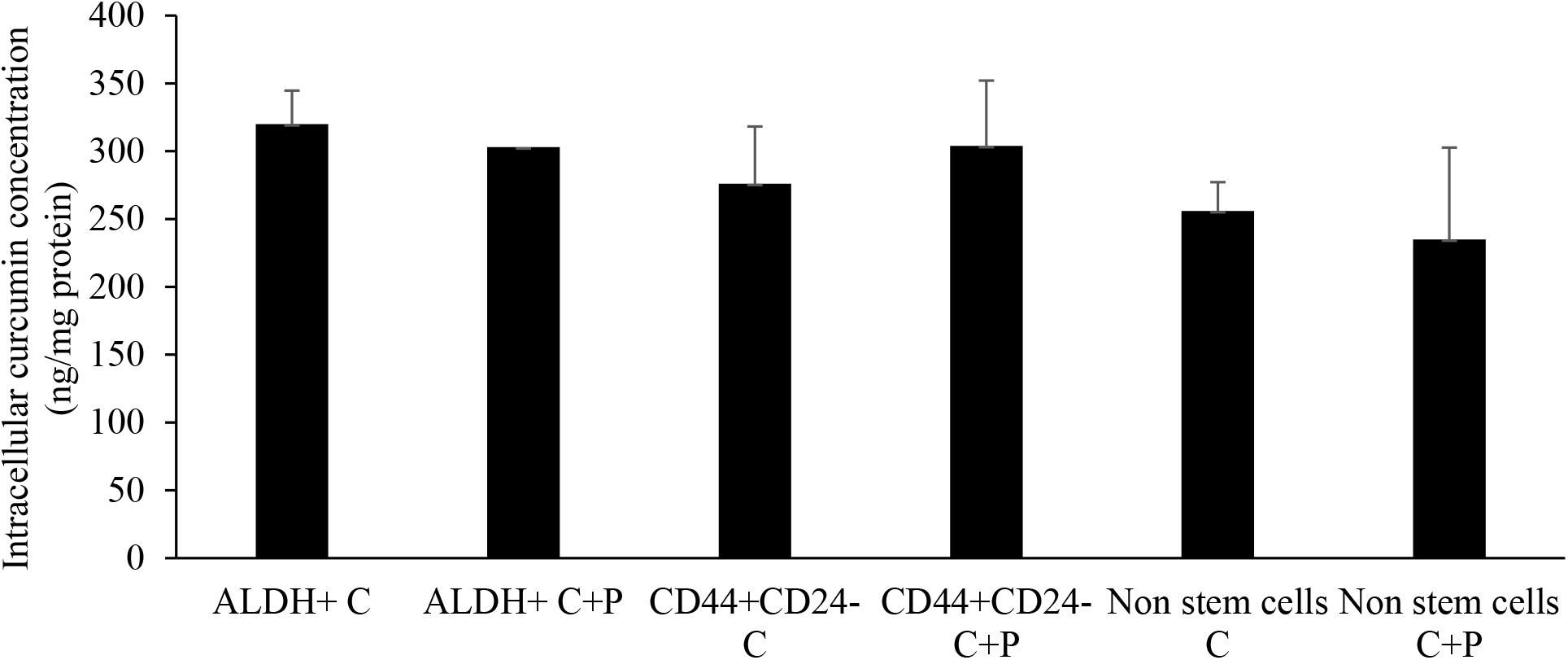
Intracellular concentrations of curcumin in ALDH^+^, ALDH^-^, CD44^+^CD24^-^ cells isolated from SUM 149 cells. After incubating ALDH^+^, ALDH^-^, CD44^+^CD24^-^, or the remaining SUM149 cells that did not express stemness cell surface markers (non-stem cells) with 15 μM curcumin ± 10 μM piperine for one hour the concentrations of curcumin were not affected by piperine co-incubation. C=curcumin, C+P= curcumin + piperine.

### Curcumin Cellular Elimination

We incubated MCF10A and SUM149 cells with curcumin 15 μM ± 10 μM piperine for 1 hour, washed and reincubated with fresh media for varying time periods from 5 min to 4 hours. We found no difference in curcumin intracellular concentration or in associated concentrations of curcumin in the fresh media of the cells incubated with or without piperine (Figure 5A). The progressive reduction of curcumin concentration intracellularly is associated with rapid increases of curcumin in the extracellular media. We also detected curcumin metabolites, tetrahydrocurcumin and curcumin sulfate conjugate in very low concentrations in the extra-cellular media (Supplemental Figure 3). The predominant form of curcumin elimination from the cell is the parent compound, curcumin. Intracellular metabolism in neoplastic cells is minimal. Piperine had no effect upon curcumin elimination, either as parent compound found in the media (Figure 5B) or in the form of metabolites in the media (Supplemental Figure 3).

**Figure 5.**
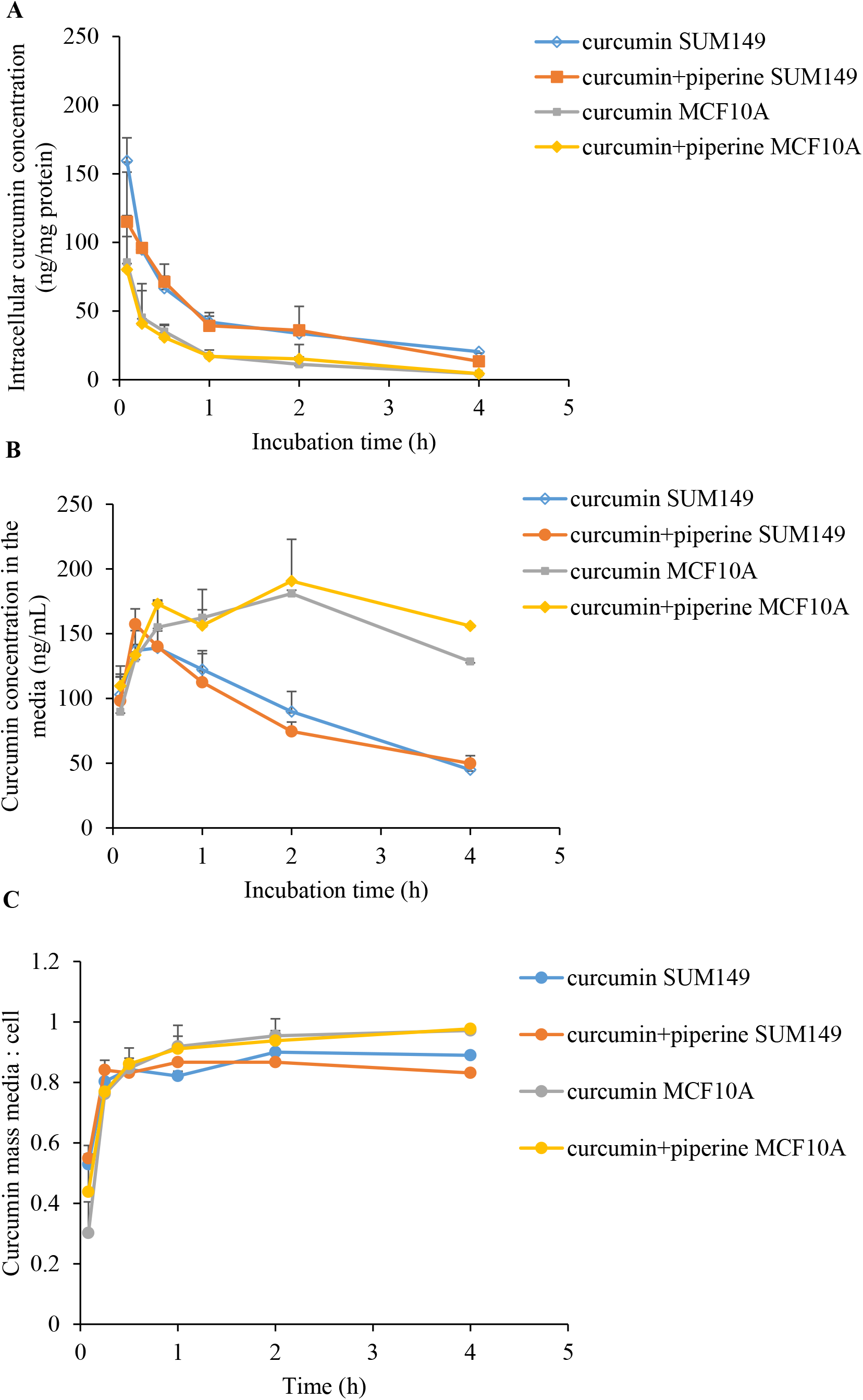
Curcumin elimination after incubation with piperine. SUM149 and MCF10A cells were incubated with curcumin 15 μM ± piperine 10 μM for 1 hour. The cells were washed and reincubated in fresh media. Panel A: Intracellular concentrations of curcumin. Panel B: Concentration of curcumin in the media. Panel C: Ratio of the curcumin mass detected by the LC-MS system in the media to the cell. Intracellular or media curcumin concentrations did not differ between the curcumin alone and the curcumin + piperine incubation groups.

## Discussion

Ayurvedic practitioners in India administer curcumin after processing it either by mixing it with other herbs or by using it in a crude turmeric preparation that consists of oils, other curcuminoids such as demethoxycurcumin, bisdemethoxycurcumin, tetrahydrocurcumin (42), and many other naturally occurring chemical excipients. The common folk-medicine practice of mixing turmeric and pepper in warm milk for treatment of respiratory or systemic inflammation, trikatu in Indian folk medicine triggered studies that showed that piperine enhances curcumin’s anti-inflammatory and anticarcinogenesis effects (7, 43, 44).

In this context and recognizing that improved analytical technologies permit a better more accurate and precise detection of curcumin and its metabolites, we assessed the cellular pharmacology of curcumin alone and in combination with piperine in a representative model of cells that mirror the carcinogenesis process. We chose cells derived from human breast neoplastic progression because of the ready availability of transformed cells (SUM149) (45), non-transformed cells (MCF10A) (46), primary normal human breast epithelial cells obtained from normal human breast mammoplasties, and cellular stemness pools that represent self-renewing stem cells (ALDH positive) (47) and quiescent stem cells (CD44^+^/CD24^-^) (47). We reproduced our prior published data demonstrating additive curcumin and piperine effects in reducing normal human mammary stem cell self-renewal and proliferation (7). While we found that piperine had similar cytotoxic effects in the MCF10A cells as the SUM149, the combination of curcumin and piperine has additive but not synergistic cytotoxic effects upon MCF10A cells. This is in contrast with the synergistic cytotoxic effect of curcumin and piperine in SUM149 cells in culture. Curcumin and piperine decreased the numbers and size of primary mammospheres derived from normal human epithelial cells. These data suggest at least an additive antiproliferative effect between curcumin and piperine.

The data we report here demonstrate that piperine has no effect on any step in the cellular uptake, metabolism, or elimination of curcumin. Piperine’s lack effect upon curcumin’s cellular absorption or elimination is consistent through the carcinogenesis continuum—transformed invasive cells represented by SUM149, non-invasive neoplastic cells represented by MCF10A, and normal human breast epithelial cells in primary culture. in the models we tested.

The higher concentrations of curcumin in the media relative to its intracellular concentrations after reincubating the cells with curcumin free media suggest that curcumin is actively transported out of the cells. Berginc, et al found that piperine significantly decreased the absorbed fraction of curcumin in Caco-2 cells and in isolated rat intestine, explained on the basis of induction of the BCRP transporter enhancing curcumin efflux from cells (29). Berginc et al use of very high concentrations of both drugs, 100 μM curcumin ± 100 μM piperine, which may not accurately model the much lower curcumin and piperine concentrations that are physiologically relevant. High concentrations of curcumin and piperine might be engaging an otherwise low affinity membrane efflux system. Although pre-administration of piperine increased curcumin bioavailability in rats in vivo (28), preincubating cells with piperine did not increase curcumin’s intracellular concentration.

We found that breast epithelial cells across the carcinogenesis continuum reduce curcumin to tetrahydrocurcumin and conjugate curcumin via sulfation. Piperine has no effects upon curcumin cellular reduction or sulfation. We did not explore piperine’s effects on curcumin glucuronidation, since curcumin is not glucuronidated in breast cells. Piperine potently inhibits glucuronidation activity in rodent models (23). In human models, 50 μM piperine did not inhibit glucuronidation activity (26, 48). Although 20 μM piperine did not inhibit the glucuronidation of Epigallocatechin gallate (EGCG) in human intestinal adenocarcinoma cells (49), piperine increased the bioavailability of the EGCG in mice (49).

An important limitation of our study is that breast cells may not serve as an adequate model in assessing epithelial cell absorption or metabolism. The absorption and conjugation of curcumin is thought to be primarily an intestinal epithelial function or hepatic function. While recognizing the limitation of not using GI epithelial cellular models, we prioritized interrogating the carcinogenesis continuum at a key clinically targeted cell for breast epithelial carcinogenesis—the breast epithelial cells. Breast epithelial cell models are readily available. Since the therapeutic context of both curcumin and piperine’s development are as potential breast cancer preventive agents, we considered cell models representative of the desired clinical target. Furthermore, readily available cellular carcinogenesis models that span the continuum of stemness to non-invasive and invasive neoplasms are not readily available for the human GI tract.

Based upon the data we present, we infer that the synergistic cytotoxic effects and inhibition of stem cell self-renewal of piperine with curcumin are due to their individual anti-carcinogenic effect. Piperine reduces size and numbers of primary mammospheres derived from normal breast epithelial cells (antiproliferative effect), and induces cytotoxicity in transformed breast neoplastic cells, non-tumorigenic neoplastic epithelial cells. We speculate that piperine’s anti-carcinogenic activity combined with curcumin may be based upon its inhibition with curcumin of one or more key stem cell control pathways such as Wnt (7, 32), Notch (50), PI3K/AKT (51), and Hedgehog (52) or downstream control mechanisms such as AP-1 and NF-κB (31), and Stat3 (53).

## Acknowledgments and Funding sources

The authors acknowledge support from The Educational Affairs and Mission sector, Ministry of Higher Education of Egyptian government, the University of Michigan’s Rogel Cancer Center Cancer Prevention Fund, the Kutche Family Memorial Chair, and the Moshe Talpaz Professorship in Translational Oncology. Support for JAC was provided by the National Institutes of Environmental Health Sciences (R01 ES028802 and P30ES01885).

