## Supplemental Figures for "Cellular Pharmacology of Curcumin With and Without Piperine"

**Supplemental Figure 1**

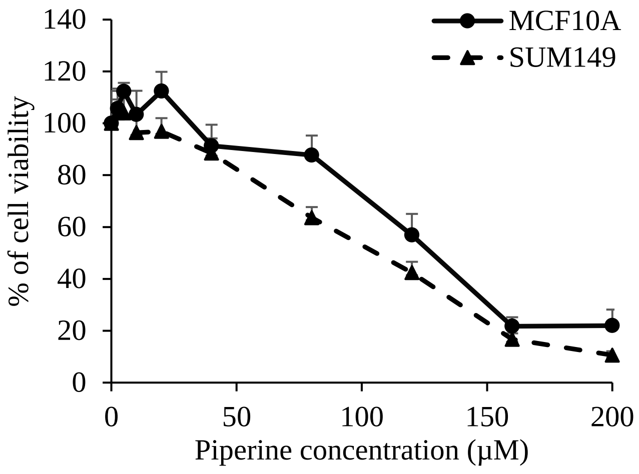


**Supplemental Figure 1.** Percent of viability of MCF10A and SUM149 cells after treating them with escalating doses of piperine (0-200 µM) compared to treatment with DMSO vehicle (0.1%) for 72 hours. Results are expressed as the percent of cell viability of compound-treated cells to vehicle-treated cells, expressed as the mean ± SD of two independent experiments 6 replicates each.

**Supplemental Figure 2**

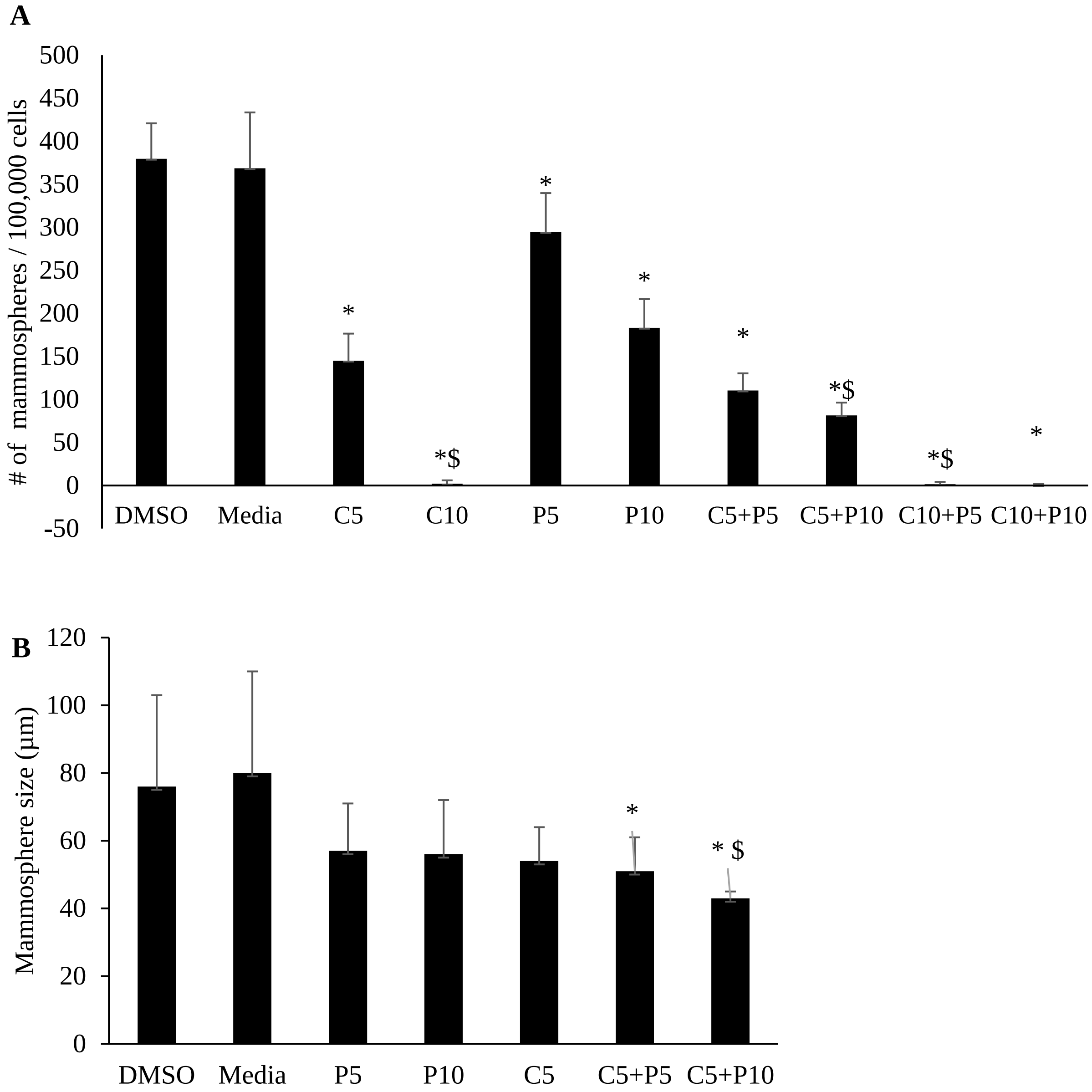


**Supplemental Figure 2. Curcumin and piperine reduce primary human breast mammosphere formation.** Number (A) and size (B) of primary mammospheres formed after treating single normal human breast cells with curcumin, piperine or both. Results are expressed as mean ± SD from three independent experiments three replicates each. Curcumin alone reduces mammosphere numbers in a dose responsive manner (C5 vs DMSO $p<0.001$; C10 vs DMSO

$p<0.0001$). Curcumin also reduces mammosphere size, but significant only at the C10 concentration ($p<0.0001$). Piperine reduces mammosphere numbers at low (P5 or P10 vs DMSO, $p<0.05$) but has no effect upon mammosphere size. The combination of curcumin and piperine at all concentrations reduces mammosphere numbers with at least an additive effect ($p<0.0001$). The effect of curcumin and piperine in combination on the reduction of mammosphere size is less potent than mammosphere numbers but statistically significant (C5+P5 $p<0.05$; C5+P10, $p<0.001$; C10+P5 or C10+P10, $p<0.0001$); *= significant from DMSO and $= significant from C5. C5, C10= curcumin 5and 10 µM, respectively. P5, P10= piperine 5and 10 µM, respectively.

**Supplemental Figure 3**

A

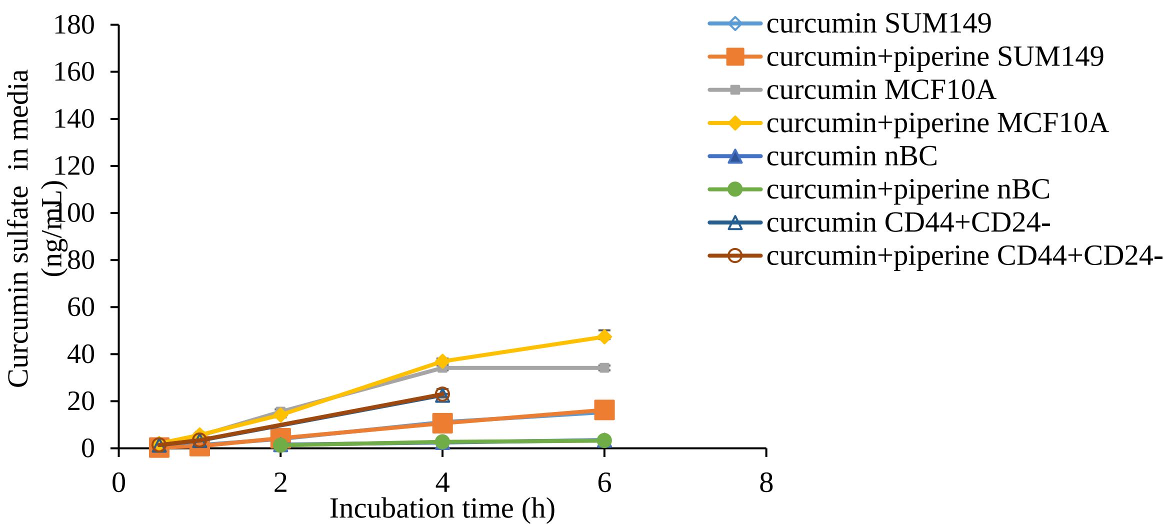


B

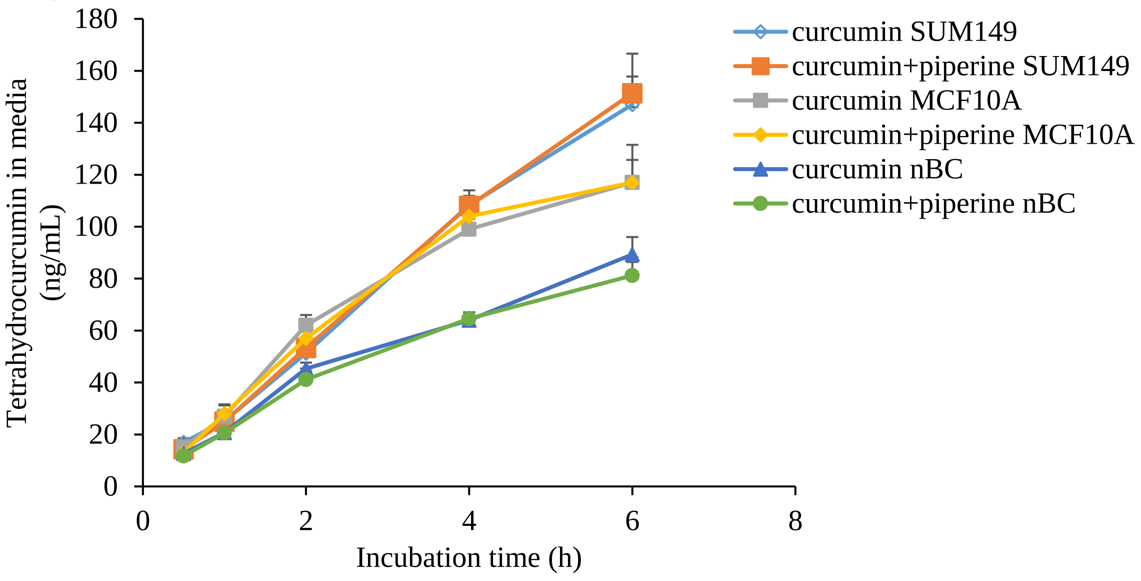


**Supplemental Figure 3. Curcumin metabolites in media.** Low concentrations of curcumin metabolites appear in media rapidly after completion of incubation with curcumin. Panel A: Curcumin sulfate conjugate concentrations versus time detected in media after incubation with 15 μM curcumin ± 10 μM piperine. Panel B: Tetrahydrocurcumin concentrations versus time.
